# Bacteriophage resistance alters antibiotic mediated intestinal expansion of enterococci

**DOI:** 10.1101/531442

**Authors:** Anushila Chatterjee, Cydney N. Johnson, Phat Luong, Karthik Hullahalli, Sara W. McBride, Alyxandria M. Schubert, Kelli L. Palmer, Paul E. Carlson, Breck A. Duerkop

**Author notes:** Correspondence: Breck A. Duerkop. A.C. and C.N.J. contributed equally to this work.

## Abstract

*Enterococcus faecalis* is a human intestinal pathobiont with intrinsic and acquired resistance to many antibiotics, including vancomycin. Nature provides a diverse and virtually untapped repertoire of bacterial viruses, or bacteriophages (phages), that could be harnessed to combat multi-drug resistant enterococcal infections. Bacterial phage resistance represents a potential barrier to the implementation of phage therapy, emphasizing the importance of investigating the molecular mechanisms underlying the emergence of phage resistance. Using a cohort of 19 environmental lytic phages with tropism against *E. faecalis*, we found that these phages require the enterococcal polysaccharide antigen (Epa) for productive infection. Epa is a surface-exposed heteroglycan synthesized by enzymes encoded by both conserved and strain specific genes. We discovered that exposure to phage selective pressure favors mutation in non-conserved *epa* genes both in culture and in a mouse model of intestinal colonization. Despite gaining phage resistance, *epa* mutant strains exhibited a loss of resistance to the cell wall targeting antibiotics, vancomycin and daptomycin. Finally, we show that an *E. faecalis epa* mutant strain is deficient in intestinal colonization, cannot expand its population upon antibiotic-driven intestinal dysbiosis and fails to be efficiently transmitted to juvenile mice following birth. This study demonstrates that phage therapy could be used in combination with antibiotics to target enterococci within a dysbiotic microbiota. Enterococci that evade phage therapy by developing resistance may be less fit at colonizing the intestine and sensitized to vancomycin preventing their overgrowth during antibiotic treatment.

## Importance

With the continued rise of multidrug resistant bacteria, it is imperative that new therapeutic options are explored. Bacteriophages (phages) hold promise for the amelioration of enterococcal infections, however, the mechanisms used by enterococci to subvert phage infection are understudied. Here, we demonstrate that a collection of phages require a cell surface exopolysaccharide for infection of *E. faecalis*. *E. faecalis* develops phage resistance by mutating polysaccharide biosynthesis genes at a cost, as this renders the bacterium more susceptible to cell wall targeting antibiotics. *E. faecalis* phage resistant isolates are also less fit at colonizing the intestine and these mutations mitigate *E. faecalis* intestinal expansion upon antibiotic selection. This study suggests that the emergence of phage resistance may not always hinder the efficacy of phage therapy and that the use of phages may sensitize bacteria to antibiotics. This could serve as a promising avenue for phage-antibiotic combination therapies.

## Introduction

Enterococci are Gram-positive commensal bacteria native to the intestinal tracts of animals, including humans (1). Under healthy conditions, enterococci exist as minority members of the microbiota in asymptomatic association with their host. However, upon antibiotic disruption of the intestinal bacterial community, enterococcal populations can flourish resulting in elevated intestinal colonization (2, 3). As dominant members of the intestinal microbiota, enterococci can breach the intestinal barrier leading to bloodstream infections (3). The pathogenic success of the enterococci is largely attributed to the development of multidrug resistance (MDR) traits, including the emergence of vancomycin resistant enterococci (VRE). *Enterococcus faecalis* and *Enterococcus faecium* represent the species most commonly associated with vancomycin resistance. In the hospital, MDR enterococcal strains can be transmitted rapidly, leading to dangerous outbreaks that put immunocompromised patients at risk (4, 5). This is especially troubling as clinical VRE isolates that are resistant to recently introduced “last-line-of-defense” antibiotics have been discovered (6-10). With limited treatment options to combat the continuing rise of MDR enterococci, it is imperative to develop alternative therapeutic approaches in addition to conventional antibiotic therapy.

Bacteriophages (phages), viruses that infect bacteria, could be used for the eradication of difficult to treat *E. faecalis* and *E. faecium* infections. Many of these are obligate lytic phages belonging to the *Siphoviridae* and *Myoviridae* families of tailed double stranded DNA phages (11). Current efforts in the development of phages as anti-enterococcal agents have focused on the treatment of systemic infections or surface associated biofilms (12-14), though they may also be effective in decolonizing the intestines of individuals in a hospital setting.

The utility of phages as effective anti-enterococcal therapeutics relies on having a detailed understanding of phage infection mechanisms and how enterococci subvert phage infection through the development of resistance. To date, only a single membrane protein, PIP_EF_ (phage infection protein of *E. faecalis*), has been definitively identified as a phage receptor for *E. faecalis* (15). The enterococcal polysaccharide antigen (Epa) is involved in phage adsorption to *E. faecalis* cells and may act as a phage receptor (16-18). Considering at least a dozen well-characterized lytic enterococcal phages have the potential to be used for phage therapy (11), identifying receptors used by phages could allow for the generation of more efficient phage cocktails to be used for the treatment of enterococcal infections. This is particularly important since phage therapies can employ the use of multivalent phage cocktails to limit the emergence of bacterial resistance (19, 20) and knowledge of phage receptors can lead to rational design of such cocktails. In addition, phages often target conserved components of the bacterial cell surface, which bacteria can mutate to subvert phage infection. If a phage receptor is essential to bacterial physiology, mutation often imposes a fitness cost (21-23). Therefore, therapeutic phages could be selected that force bacterial targets to trade a fitness benefit in return for phage resistance, making them less pathogenic and possibly more susceptible to current antimicrobials (21, 24).

The *E. faecalis* genome contains both broadly conserved and strain variable *epa* genes (25). Using *E. faecalis* and a collection of uncharacterized virulent phages, we identify genes located in the variable region of the *epa* locus to be critical for phage infection. Epa is directly involved in phage attachment to the bacterial surface. Exposure of *E. faecalis* to certain phages, both *in vitro* and in the mouse intestine, selects for mutations in the *epa* locus, primarily in *epa* variable genes. Loss of function mutations in two *epa* variable genes, *epaS* and *epaAC*, resulted in cell surface alterations that increase the sensitivity of *E. faecalis* to cell wall targeting antibiotics. During colonization of the mouse intestine, an *E. faecalis epaS* mutant had a colonization defect in both adult mice and juvenile mice shortly following birth. The *epa*S mutant also failed to efficiently outgrow in the intestine upon antibiotic mediated perturbation of the native commensal bacteria. Together these data suggest that during enterococcal intestinal dysbiosis, phages could be harnessed to selectively modify the enterococcal population in favor of *epa* mutants that could be targeted more efficiently with concurrent antibiotic therapies.

## Results

### Host range and morphology of enterococcal bacteriophages

We obtained a library of 19 enterococcal specific bacteriophages through the Biological Defense Research Directorate of the Naval Medical Research Center (NMRC). These phages were isolated from environmental sources as described previously (26). Phage spot-agar assays were performed (27, 28) to assess phage infectivity against 21 *E. faecalis* strains whose susceptibility profiles for these phages were unknown. Phage lysates formed clear, opaque or no spots against specific *E. faecalis* strains, indicating strong infection, weak infection or no infection, respectively (Fig. 1A). With the exception of phi44 and phi49, each phage infected at least one *E. faecalis* strain and there was host range variability among the phages. Phages phi4, phi17, and phi19 had the broadest host range, infecting more than 75% of the *E. faecalis* strains tested. In contrast, phages phi16, phi35, phi47, phi48, and phi51 had restricted host ranges, infecting four or less *E. faecalis* strains (Fig. 1A). Hence, this phage collection includes both broad and narrow host range phages.

**Figure 1.**
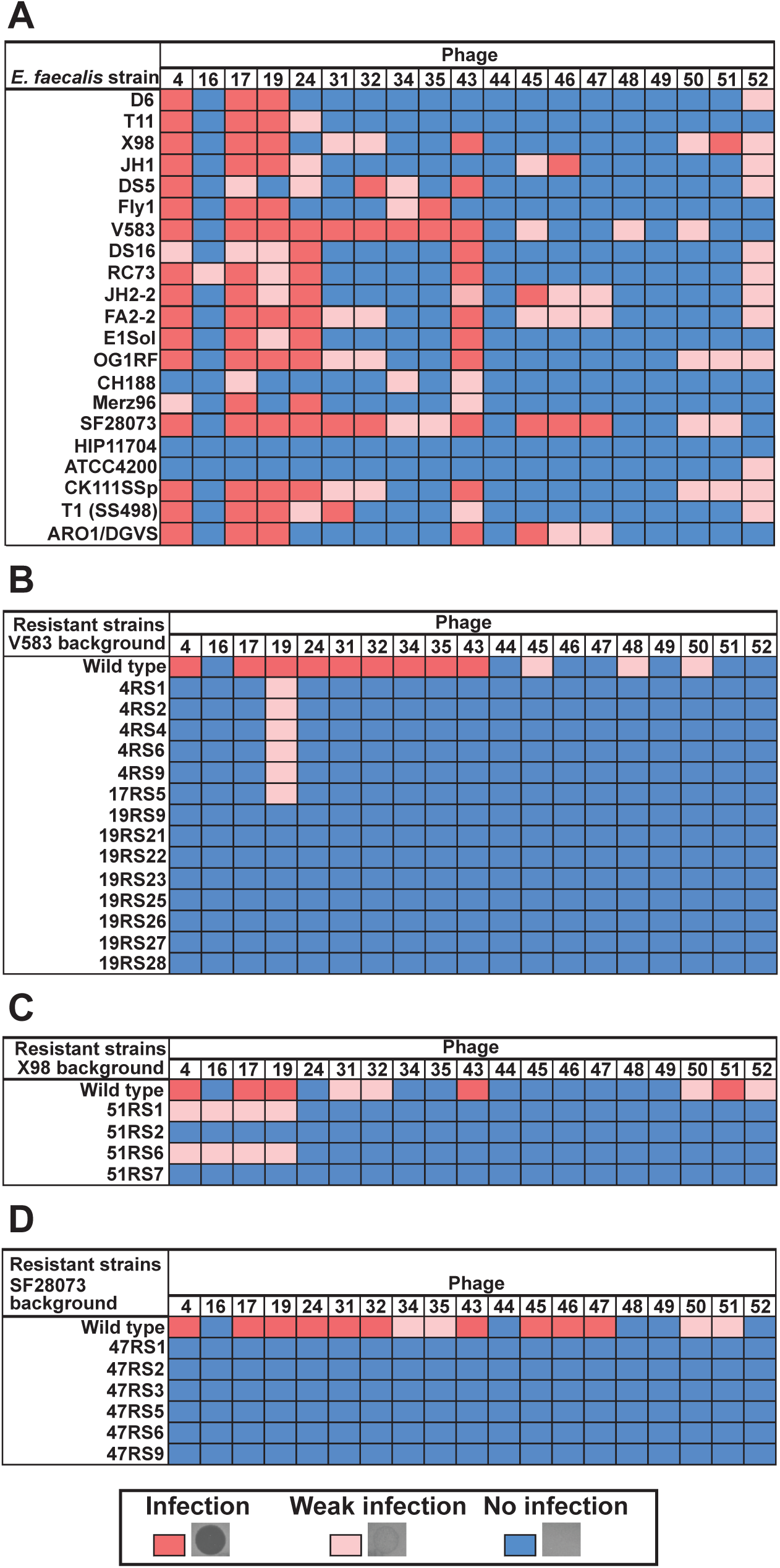
NMRC phages have broad and narrow *E. faecalis* host ranges. Red, pink and blue boxes represent the results of phage infection spot assays indicating infection, weak infection and no infection, respectively. **(A)** The host range of 19 NMRC phages against 21 different *E. faecalis* strains. **(B – D)** Phage sensitivity profiles of *E. faecalis* phage resistant isolates indicates a high degree of cross infectivity among NMRC phages. **(B)** phi4, phi17 and phi19 resistant strains have lost susceptibility to the majority of phages that can infect the V583 parental strain. **(C)** Compared to the wild type X98, phi51 resistant mutants gained immunity against all phages capable of infecting X98. **(D)** Spot assays demonstrate that phi47 resistant *E. faecalis* are now resistant to phages phi4, phi17 and phi19.

We performed transmission electron microscopy to determine the structural features of three broad host range and two narrow host range NMRC phages. Phages phi4, phi47 and phi51 belong to the *Siphoviridae* family of long non-contractile tailed phages. phi4 has a cubic icosahedral capsid symmetry (Fig. 2A), whereas phi47 and phi51 have elongated prolate capsids (Fig. 2B and 2E) (29). Phages phi17 and phi19 belong to the *Myoviridae* family with icosahedral capsids and sheathed contractile tails (Fig. 2C and 2D) (29).

**Figure 2.**
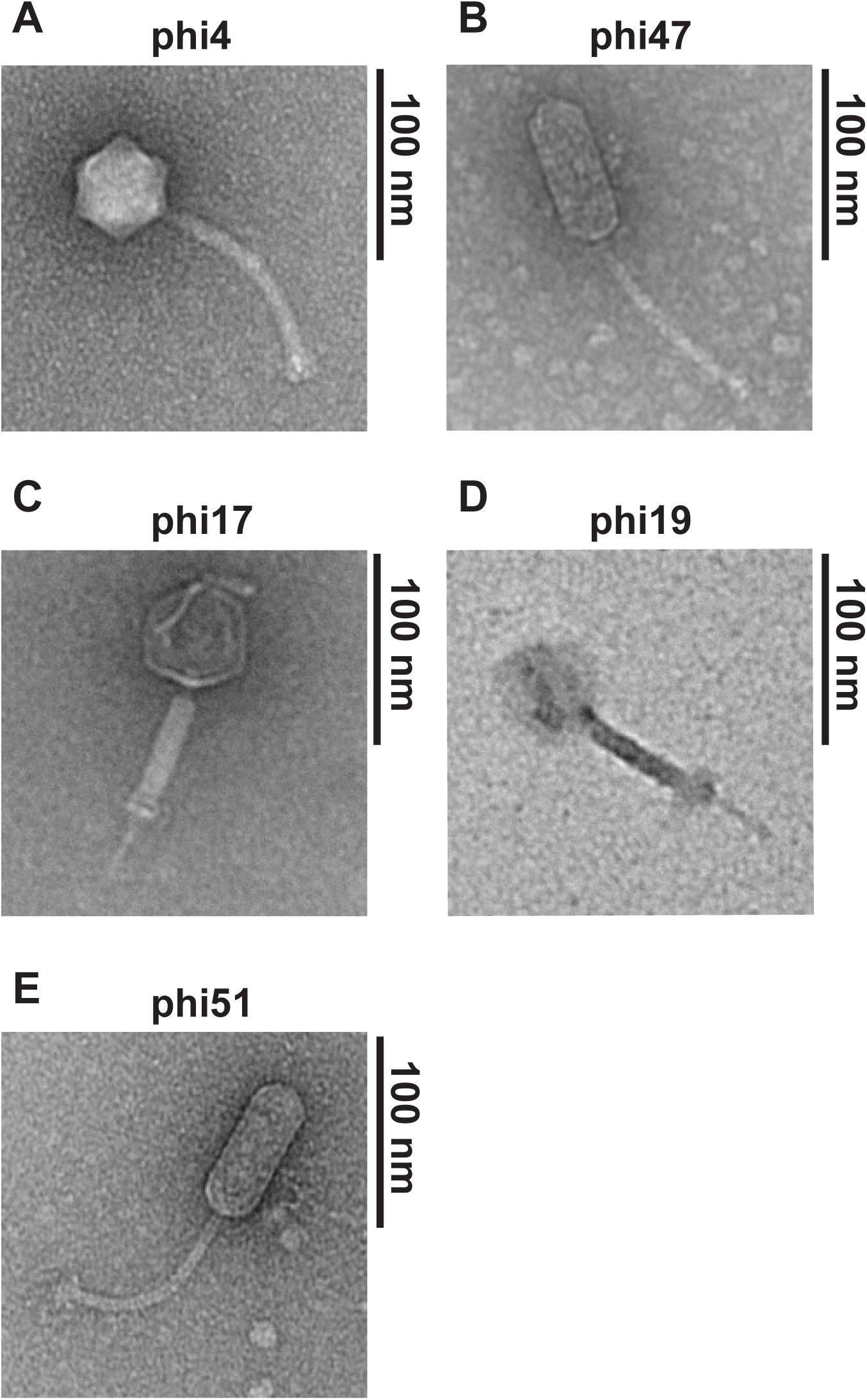
Transmission electron microscopy of NMRC phages reveals diverse morphologies. All phages imaged are double stranded DNA phages of the order Caudovirales. Phages phi4 **(A)**, phi47 **(B)** and phi51 **(E)** are *Siphoviridae*. Phage phi17 **(C)** and phi19 **(D)** are *Myoviridae*.

### NMRC phages infect E. faecalis independent of PIP_EF_

To determine how NMRC phages infect *E. faecalis*, we tested their ability to infect *E. Faecalis* BDU50, a *pip*_*EF*_ mutant strain of *E. faecalis* V583 that is resistant to phage infection (15). Phages from the NMRC collection had identical tropism for both wild type *E. faecalis* V583 and BDU50, indicating that NMRC phages infect *E. faecalis* in a PIP_EF_-independent manner (Fig. 3A and 3B). To determine the molecular mechanism underlying NMRC phage infection, we selected *E. faecalis* phage resistant isolates using both broad (phi4, phi17, phi19) and narrow (phi47, phi51) host range phages, using *E. faecalis* strains V583 (phi4, phi17, phi19), SF28073 (phi47) and X98 (phi51) (Table S1). Phages were mixed with *E. faecalis* in top agar, poured over the surface of an agar plate, and assayed for confluent lysis. Potential phage resistant colonies emerged within zones of lysis after overnight incubation. The spot-agar assay was used to confirm phage resistance of the isolates (Fig. 1B-D).

**Figure 3.**
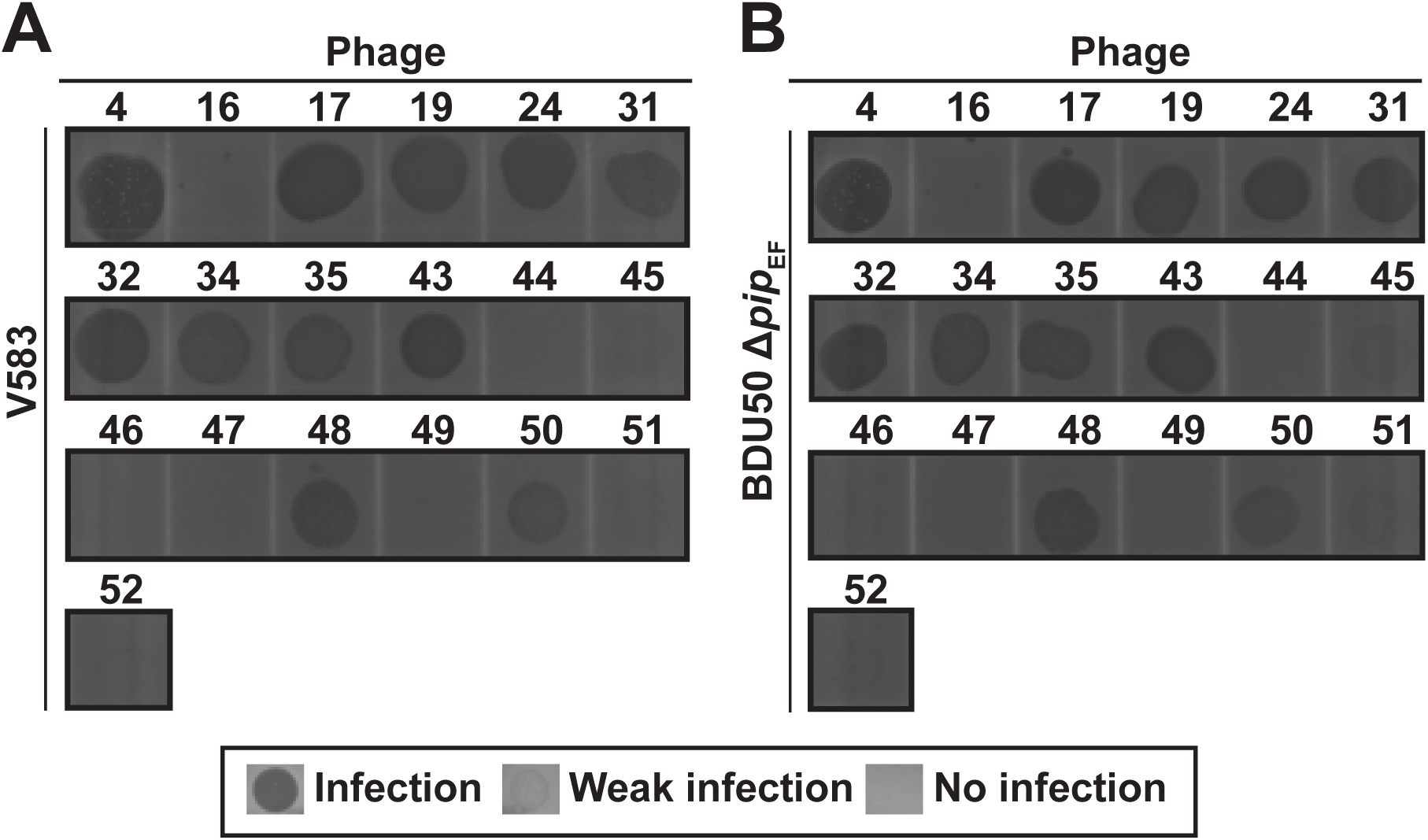
NMRC phages infect *E. faecalis* independent of PIP_EF_. NMRC phages were spotted onto **(A)** wild type *E. faecalis* V583 or **(B)** *E. faecalis* BDU50 a *Δpip_EF_* isogenic mutant of strain V583.

We next determined the extent of phage cross-resistance by testing the *E. faecalis* phage resistant isolates against all other phages in the NMRC collection (Fig. 1B-D). Our data show that regardless of the phage used to select for resistance, there is broad cross-resistance to other phages from the collection (Fig. 1B-D). These data suggest that even though NMRC phages have distinct host tropisms they likely infect through a related mechanism.

### Mutations in enterococcal polysaccharide antigen (epa) genes promote phage resistance

To determine the genetic basis of PIP_EF_-independent lytic phage infection in *E. faecalis*, we performed whole genome sequencing of select spontaneous phage resistant mutants and their corresponding wild type parental strains. In all cases, no matter which phage was used to select for resistance, phage resistant isolates harbored mutations in the enterococcal polysaccharide antigen (*epa*) gene cluster (Table 1, Fig. 4A), implicating *epa* mutation as a key contributor to phage resistance.

**Figure 4.**
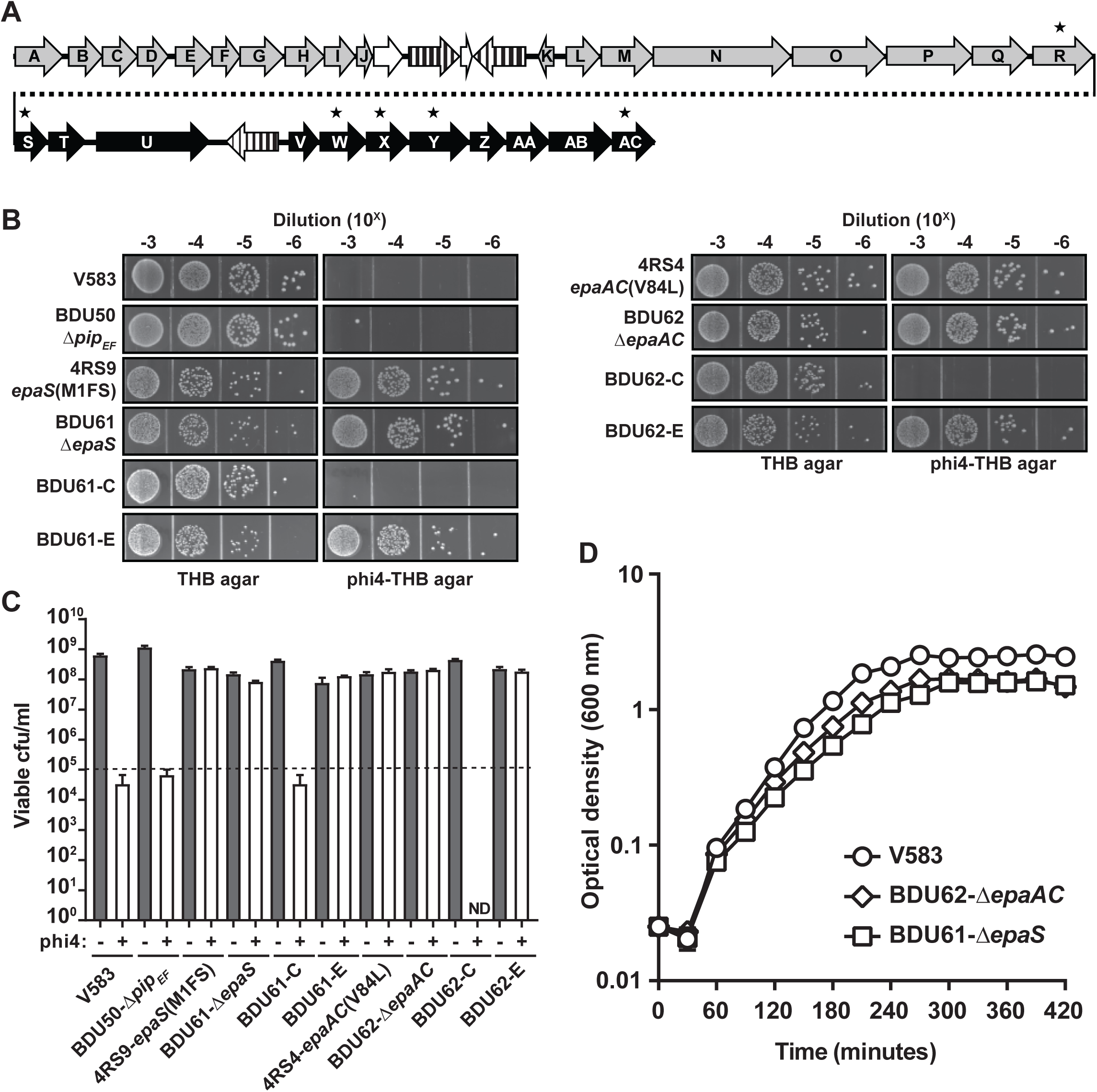
Mutations in the *epa* locus confer phage resistance. **(A)** Schematic depicting the *epa* locus of *E. faecalis* V583. The core *epa* genes (*epaA* – *epaR*) found in all *E. faecalis* strains are shown in grey. *epa* variable genes (*epaS* – *epaAC*) downstream of the conserved core genes are shown in black. Genes with vertical black lines indicate insertion sequence elements IS256 or ISEf1. White genes indicate a putative racemase gene that has been disrupted by an IS256 element. Stars designate genes where mutations were found in sequenced phage resistant isolates. All the genes are drawn to scale. **(B – C)** phi4 susceptibility assays were performed on serially diluted overnight cultures of specific bacterial strains. Dilutions were spotted onto THB agar plates with or without 57×10^8^ pfu/ml of phi4. Representative spot plates **(B)** and the corresponding quantitative viable colony counts **(C)** are shown. **(D)** Growth curves comparing wild type *E. faecalis* V583 and isogenic *epa* mutant strains in BHI broth. The dashed horizontal line in C indicates the limit of detection based on the bacterial plating procedure. -E (empty vector) and - C (complemented). Open bars below the limit of detection in **(C)** occur when one or more colonies arise for a single experimental replicate at the 10^-3^ dilution. ND – none detected.

The *epa* locus encodes genes involved in the biosynthesis of a rhamnose containing cell surface associated polysaccharide (30), yet the biochemical functions of most Epa proteins are uncharacterized. The *E. faecalis epa* gene cluster (Fig. 4A) consists of a conserved core set of 18 genes (*epaA* – *epaR*) upstream a group of variable genes beginning at *epaS* (EF2176 in V583) and ending at *epaAC* (EF2165 in V583) (17, 25). *epaR*, encoding a transmembrane glycosyltransferase, was the only core gene found to be mutated (Table S1). This is consistent with a recent study demonstrating that mutation of *epaR* in *E. faecalis* OG1RF results in resistance to infection by the phage NPV1 (16). The remaining mutations in the phage resistant isolates mapped to variable region *epa* genes including *epaS, epaW, epaX* and *epaAC* (Fig. 4A, 4B, and Table S1). Notably, *epaX* was recently found to aid in the adsorption of the *Podoviridae* phage Idefix (18).

Since *E. faecalis* is a native inhabitant of the intestine, we determined whether phage driven *epa* mutations arose in germ free C57BL6/J mice colonized with the *E. faecalis* strains V583 or SF28073. Mice were treated orally with 10^10^ pfu of phi4 or phi47 for seven days, after which bacteria were isolated from the feces and screened for phage resistance by plating on agar plates containing either phi4 or phi47. We sequenced six *E. faecalis* V583 isolates resistant to phi4 and ten *E. faecalis* SF28073 isolates resistant to phi47. Similar to the phage resistant isolates acquired *in vitro*, all *in vivo* phage resistant isolates had mutations that mapped to the *epa* locus (Table S2). Mutations were restricted to *epaR* and *epaS* in the *E. faecalis* SF28073 resistant isolates. Two of the six *E. faecalis* phi4 resistant isolates in the V583 background had mutations that mapped to *epaX* and *epaAC*. The remaining four isolates had mutations that mapped to *epaY* (Table S2). Interestingly, no *epaY* mutations were found in any of 16 *in vitro* derived *epa* specific phage resistant isolates (Table S1), suggesting that *in vivo* phage selective pressure may be directed toward alternative *epa* variable genes.

Previous studies have demonstrated that core *epa* genes are important for phage infection of *E. faecalis* (16-18). To confirm the role of *epa* variable genes in facilitating phage infection of the NMRC phages, we generated in-frame deletion mutants of *epaS* and *epaAC* in *E. faecalis* V583 using allelic replacement. The mutants achieved a slightly lower overall culture density in the stationary phase (Fig. 4D) but had a similar doubling time during logarithmic growth compared to wild type *E. faecalis* V583 (31 min - wild type, 35 min - Δ*epaS*, 33 min - Δ*epaAC*). We attempted to make unmarked deletions in *epaX* and *epaW*, however, we were unable to generate these mutant strains. Because *epa* mutations resulted in widespread resistance of *E. faecalis* to many of the phages in the NMRC collection (Fig. 1B), we chose to confirm these isogenic mutants using phi4. As judged by bacterial growth on agar plates containing phi4, *E. faecalis* strains BDU61 (Δ*epaS*) and BDU62 (Δ*epaAC*) were phenotypically indistinguishable from the spontaneous phage resistant isolates 4RS4 and 4RS9, respectively (Fig. 4B and 4C). phi4 susceptibility could be restored by complementation (Fig. 4B and 4C). Together, these data indicate that the loss-of-function of *epaS* and *epaAC* alone are sufficient to confer phage resistance.

### Epa dictates phage adsorption but not phage infectivity of E. faecalis

To investigate how Epa contributes to phage infection, we tested the ability of phages to adsorb to phage resistant or wild type *E. faecalis* cells. phi4 adsorption was higher for wild type *E. faecalis* V583 compared to the *epa* mutant isolates 4RS4, 4RS9, 17RS5, 19RS21 and 19RS28 (Fig. 5A). Consistent with this observation, in frame deletion of *epaS* or *epaAC* abrogated the adsorption of phi4 to *E. faecalis* (Fig. 5A). These data indicate that Epa cell wall modification is essential for phage adsorption.

**Figure 5.**
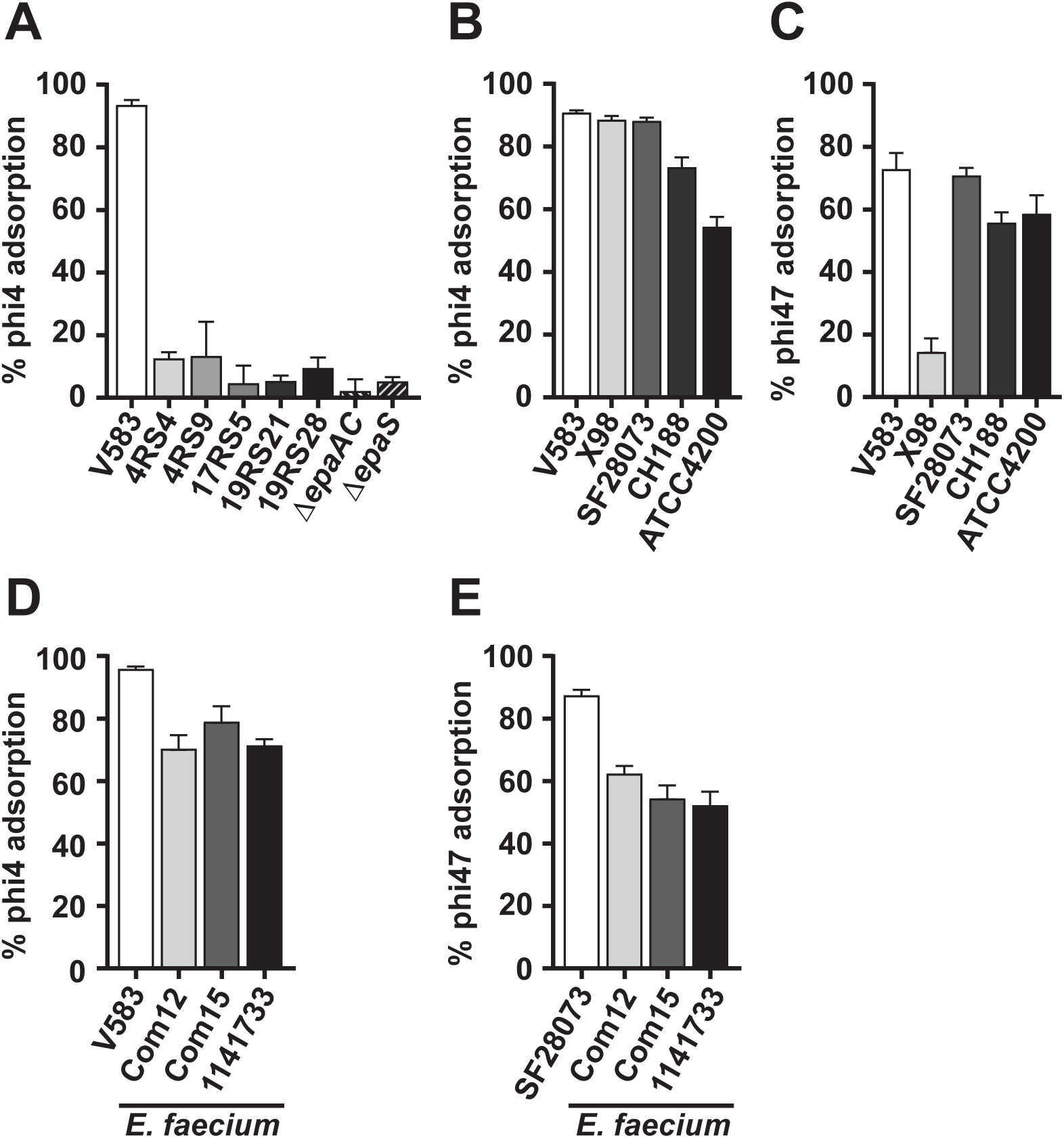
NMRC phages adsorb to a broad array of enterococci through Epa. **(A)** Wild type *E. faecalis* V583 but not spontaneous *epa* mutants or isogenic *epa* deletion strains of *E. faecalis* efficiently adsorb phi4. **(B)** phi4 adsorption profile of cognate (V583, X98 and SF28073) and non-cognate (CH188 and ATCC4200) *E. faecalis* strains. **(C)** phi47 adsorption profile of cognate (SF28073) and non-cognate (V583, X98, CH188 and ATCC4200) *E. faecalis* strains. **(D – E)** phi4 and phi47 adsorption to various strains of the related bacterium *E. faecium*.

We next assessed whether susceptibility to infection by specific phages is dictated by the ability of phages to adsorb to the surface of *E. faecalis* cells. phi4 adsorbed to both cognate (V583, X98 and SF28073) and non-cognate (CH188 and ATCC4200) *E. faecalis* cells (Fig. 5B). Similarly, phi47 adsorbed to *E. faecalis* SF28073 and adsorbed with ∼60-80% efficiency to strains that it does not infect (V583, CH188 and ATCC4200) (Fig. 5C and Fig. 1A). To determine if the promiscuous adsorption observed for phi4 and phi47 was restricted to *E. faecalis*, we tested the ability of these phages to adsorb to the related enterococcal species *E. faecium*. Greater than 50% of phi4 and phi47 phage particles adsorbed to *E. faecium* strains Com12, Com15 and 1141733 (Fig. 5D – 5E). *E. faecium* harbors an *epa* locus which resembles the *epa* locus of *E. faecalis* (25). From these data we conclude that Epa is important for primary phage adsorption prior to infection. Considering phages phi4 and phi47 adsorb to strains that are naturally resistant to infection or killing, suggests that either abortive infection drives this resistance or an unidentified receptor required for DNA entry dictates phage infectivity.

### epa variable gene mutations increase susceptibility to cell wall targeting antibiotics and alter cell surface properties

Previous work showed that mutation of the genes *epaI, epaR and epaOX* (*epaX* in strain V583), increase the susceptibility of *E. faecalis* OG1RF to the cell membrane specific antibiotic daptomycin (16, 31). We sought to determine if the loss of *epa* variable genes, *epaS* and *epaAC*, conferred similar enhanced sensitivity to cell wall targeting antibiotics. To test this, we compared the sensitivity of wild type*E. faecalis* V583 and isogenic *epaS* and *epaAC* mutants to daptomycin and vancomycin. We chose to assess vancomycin sensitivity because its mechanism of action targets cell wall biosynthesis and *E. faecalis* V583 is vancomycin resistant (32). The *epaS* mutant showed increased susceptibility to both vancomycin and daptomycin compared to wild type *E. faecalis* V583 (Fig. 6A and 6B). To a lesser extent the *epaAC* mutant also showed increased susceptibility to both antibiotics albeit at concentrations higher than those observed for the *epaS* mutant (Fig. 6C and 6D). These data indicate that similar to other *epa* genes, *epaS* and *epaAC* play a role in the structural integrity of the *E. faecalis* cell wall during antibiotic pressure.

**Figure 6.**
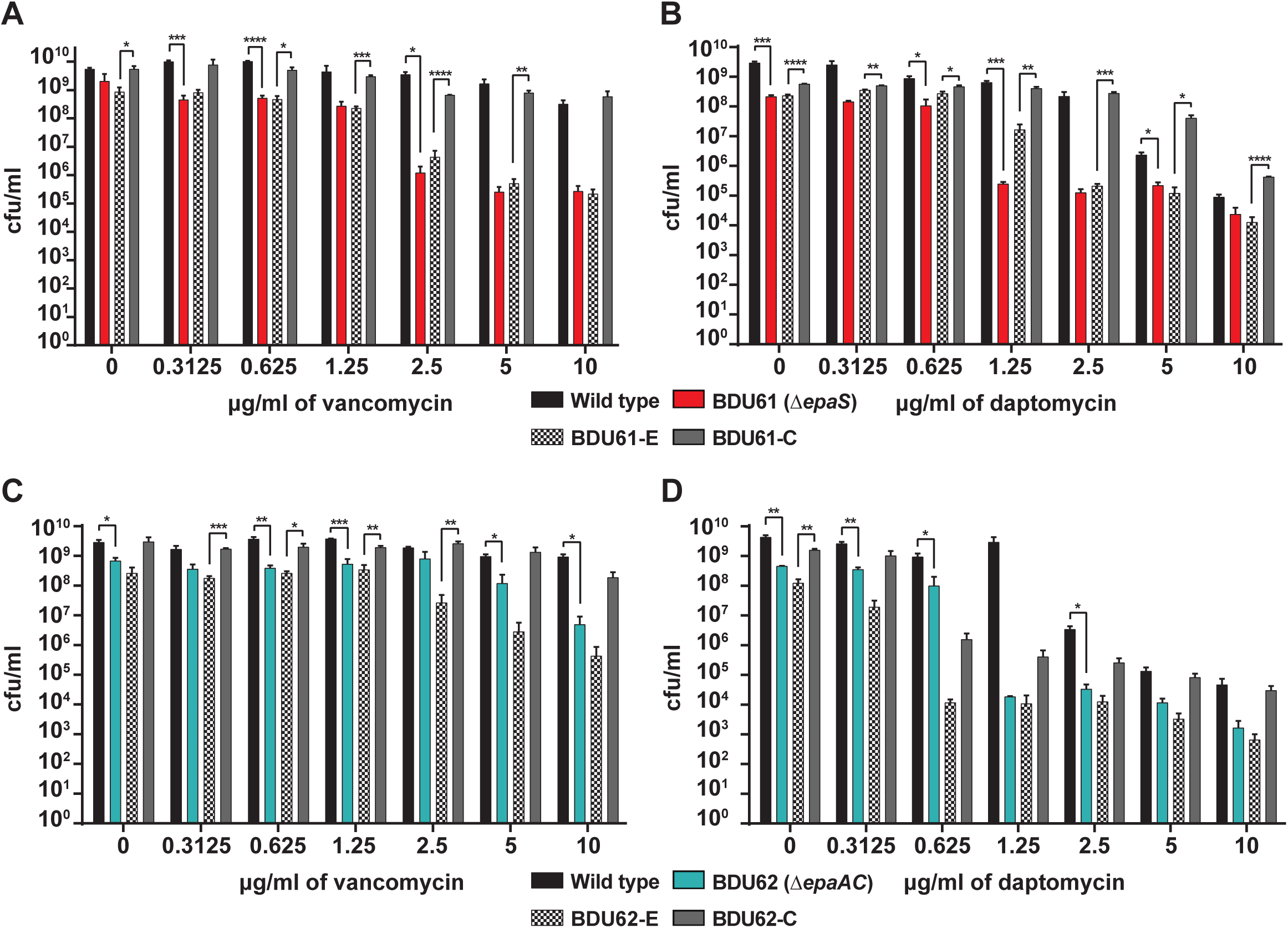
*E. faecalis epa* mutant strains are more susceptible to cell wall targeting antibiotics. Antibiotic susceptibility profiles of wild type *E. faecalis* V583 and the *epa* mutant strains BDU61 *(ΔepaS*) and BDU62 *(ΔepaAC*). Vancomycin susceptibility **(A and C)** and daptomycin susceptibility **(B and D)** of the mutants was compared to wild type *E. faecalis* V583 and complementation strains, -E (empty vector) and -C (complemented). *p<0.01, ** p<0.001, ***p<0.0008, ****p<0.0001 by Student’s t-test.

*Epa* mutations likely result in modified cell wall anchored sugar composition (16, 17). To determine if these modifications influence the overall charge of the *E. faecalis* cell wall, we performed a protein binding assay using the cationic protein cytochrome *c* (33). Cytochrome *c* bound less to the *epaS* and *epaAC* mutants compared to the wild type strain (Fig. S1). Complementation restored cytochrome *c* binding to wild type levels (Fig. S1). These data suggest that the cell wall of the *epaS* and *epaAC* mutants has a greater net positive charge compared to wild type *E. faecalis* V583, confirming that loss of function mutations in the *epa* variable genes influence cell surface charge.

### EpaS supports colonization and antibiotic mediated expansion of intestinal E. faecalis

Recently, Rigottier-Gois *et al* (34) reported that mutation of the *E. faecalis epa* variable gene *epaX,* encoding a group 2 glycosyltransferase domain protein, results in an intestinal colonization defect in mice. EpaS also contains a group 2 glycosyltransferase domain. Therefore, we tested whether an *epaS* deletion strain has an intestinal colonization defect. Groups of conventional C57BL6/J mice were colonized with either wild type *E. faecalis* V583 or an isogenic *epaS* mutant strain by oral gavage followed by addition of the bacteria to the drinking water for 18 days (Fig. 7A). On day 18 the mice were given bacteria-free water and the *E. faecalis* colonization levels were monitored for 10 days. Both the wild type and *epaS* mutant established persistent colonization over the 10 day period (Fig. 7B). However, beginning three days post removal of bacteria from the drinking water (day 21), the *epaS* mutant colonized ∼33-fold lower than wild type *E. faecalis* V583 (Fig. 7B). Thus, similar to *epaX, epaS* is a colonization factor. This also suggests that Epa glycosyltransferases are critical for intestinal colonization.

**Figure 7.**
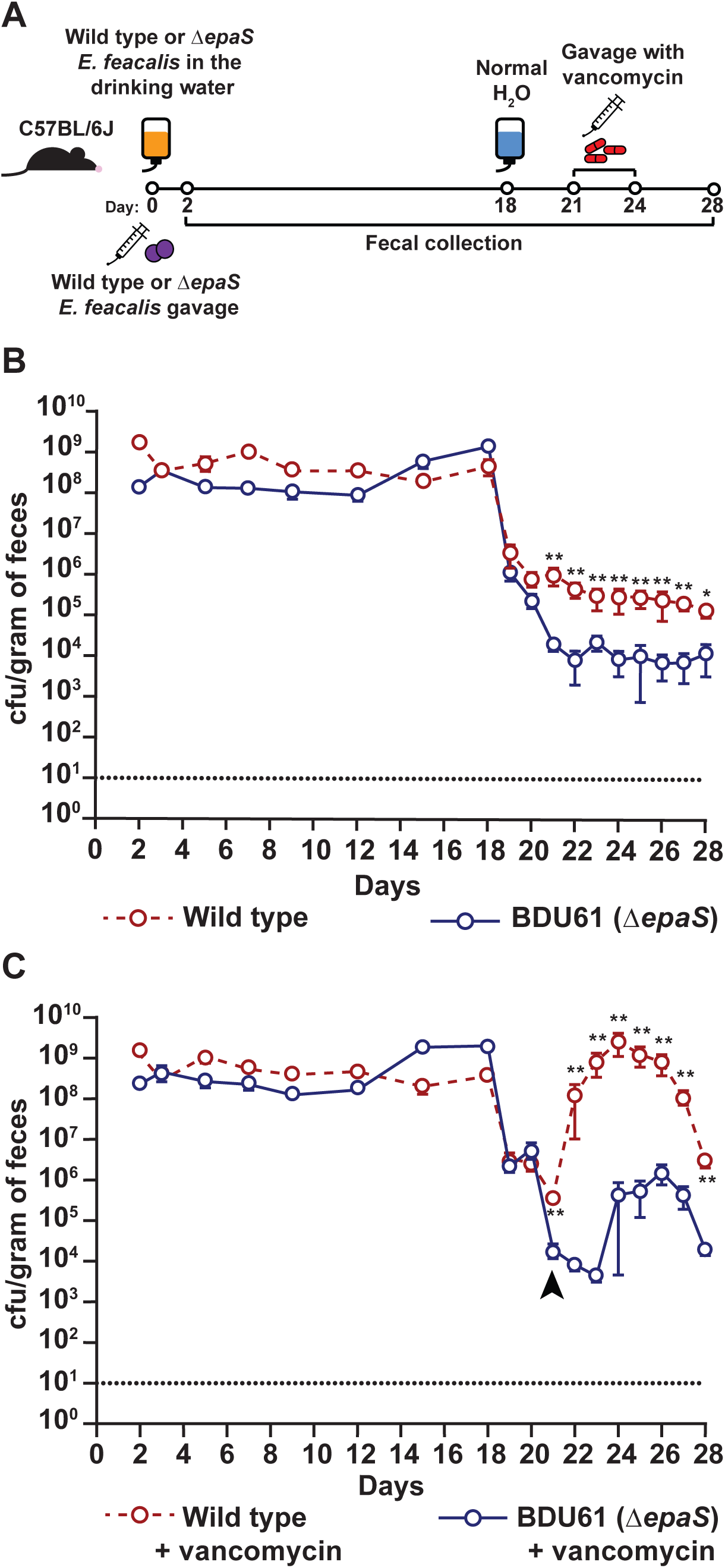
Mutation of *epaS* ameliorates antibiotic mediated expansion of intestinal *E. faecalis*. **(A)** Cartoon depicting the regiment of bacterial and antibiotic exposure to mice. **(B)** Colonization of conventional mice with either wild type *E. faecalis* V583 or the isogenic *epaS* mutant strain. **(C)** Colonization of conventional mice with either wild type *E. faecalis* V583 or the isogenic *epaS* mutant strain. At day 21 (indicated with an arrowhead), following the introduction of bacteria-free water, the mice were orally treated with 100 µg of vancomycin daily for four days. The dashed horizontal lines in **(B)** and **(C)** indicates the limit of detection based on the bacterial plating procedure. *p<0.04, **p<0.008 by Student’s t-test with Mann-Whitney U correction.

Enterococcal intestinal dysbiosis has been linked to antibiotic use in humans and these individuals are at increased risk of developing enterococcal blood stream infections (3, 35). Having observed that the *epaS* mutant is more susceptible to vancomycin treatment *in vitro* (Fig. 6A), we asked whether functional EpaS would be beneficial during antibiotic mediated *E. faecalis* expansion in the intestine. To test this, we performed an experiment identical to that described in Fig. 7B, except that starting on day 21 the mice were gavaged with 100 µg of vancomycin daily for four days (Fig. 7C). Immediately following the first dose of vancomycin, mice colonized with wild type *E. faecalis* V583 experienced a 4-log increase in *E. faecalis* colonization compared to mice colonized with the *epaS* mutant strain (Fig. 7C). During the course of vancomycin treatment, the *epaS* mutant remained at a colonization level significantly lower level than wild type *E. faecalis* V583. Three days post vancomycin treatment we observed a slight bloom of the *epaS* mutant; however, the *epaS* mutant did not achieve a similar level of colonization compared to wild type *E. faecalis* V583. We hypothesize that this bloom may be due to vancomycin mediated killing of commensal bacteria, freeing up previously occupied niches that allow the *epaS* mutant to expand its population. Considering the *epaS* mutant strain does not rebound to the same levels as wild type *E. faecalis* V583 following vancomycin treatment, these data show that antibiotic induced intestinal expansion of the enterococci requires functional EpaS.

### EpaS is required for successful transmission of E. faecalis to newborn mice

Studies suggest that offspring acquire commensal *E. faecalis* from mother’s breastmilk and the vaginal tract during birth (36). However, nothing is known about the mechanisms that contribute to the ability of antibiotic resistant *E. faecalis* to transmit to and colonize the intestine following birth. Therefore, we tested the ability of wild type *E. faecalis* V583 and the isogenic *epaS* mutant strain to be transmitted to naïve mouse pups born to mothers colonized with the bacteria. Female C57BL6/J mice were impregnated while continuously exposed to wild type *E. faecalis* V583 or the *epaS* mutant in their drinking water. After 21 days of bacterial exposure, the pregnant mothers were switched to clean water. The mothers littered their pups within 5-8 days after transitioning to bacteria-free drinking water. Mothers were chronically colonized for the duration of the experiment, however, the *epaS* mutant was maintained at a lower level relative to the wild type *E. faecalis* V583 (Fig. S2). After weaning (3 weeks post birth), we determined the levels of wild type *E. faecalis* V583 and the *epaS* mutant in the feces of the pups. Recovery of wild type *E. faecalis* V583 from the pups was significantly higher in comparison to the *epaS* mutant (Fig. 8). Therefore, our data suggest that EpaS is an important factor for the colonization and transmission of *E. faecalis* to newborns.

**Figure 8.**
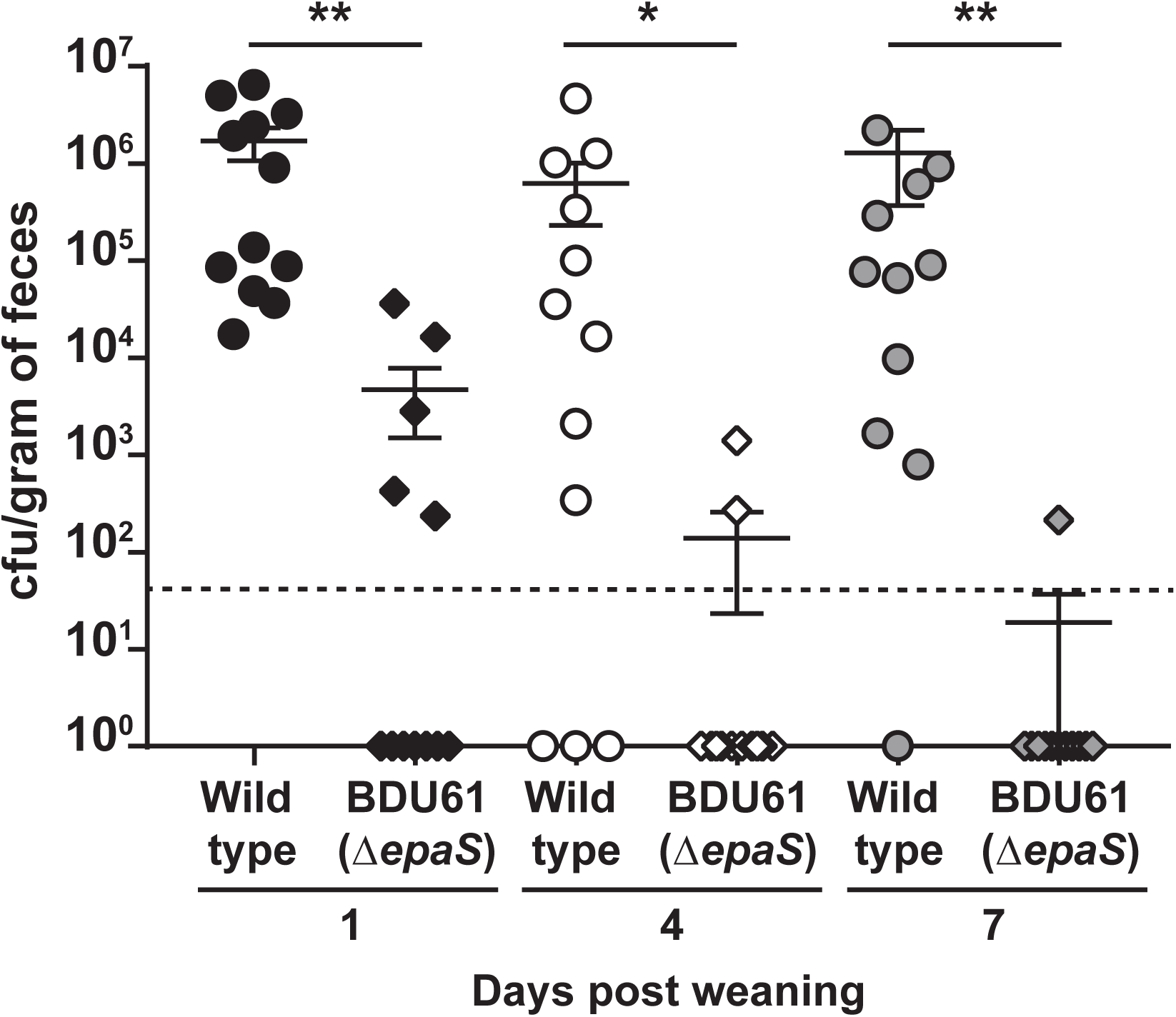
Mutation of *epaS* prevents transmission to and colonization of offspring born to chronically colonized mothers. Fecal abundance of wild type *E. faecalis* V583 and the *epaS* mutant strain BDU61 from juvenile mice born to chronically colonized mothers. Data show the cfu/gram of feces on day 1 (3 weeks after birth), day 4 and day 7 post weaning. The dashed horizontal line indicates the limit of detection based on the bacterial plating procedure. *p=0.001, **p<0.0001 by Student’s t-test with Mann-Whitney U correction.

## Discussion

Enterococci have developed and acquired resistance to antibiotics and continue to do so. Thus, there is renewed interest in the use of phages for the treatment of MDR infections. Understanding the molecular mechanisms underlying phage-host interactions could aid in the development of phage therapies by influencing the design of effective phage cocktails. In the current study, we assessed the infectivity of lytic phages that kill *E. faecalis* and demonstrated that genes in the variable region of the *epa* locus are involved in phage infection. Exposure to phages, both *in vitro* and *in vivo*, promoted the acquisition of phage resistance. Interestingly, phage resistant *E. faecalis* strains harboring loss-of-function mutations in the variable genes, *epaS* and *epaAC*, resulted in the sensitization of the bacteria to cell wall targeting antibiotics. Additionally, an *epaS* mutant was unable to efficiently colonize the mouse intestine of adult and juvenile mice in the presence of a conventional microbiota, and failed to overgrow during vancomycin treatment. This suggests that overgrowth of vancomycin-resistant *E. faecalis* in patients could be prevented using phage therapy. More broadly, our data suggest that phages could be used to exploit the evolution of bacterial phage resistance as an adjuvant to antibiotic therapy, in cases where acquisition of phage resistance leads to new antibiotic sensitivities.

Bacterial surface polysaccharides directly interact with mammalian host surfaces and are key virulence factors (17, 34, 37-41). Previous studies identified glucose, rhamnose, *N*-acetylglucosamine, *N*-acetyl galactosamine, and galactose as major components of Epa (17, 34); however, there are gaps in our understanding of the Epa cell surface architecture. Epa is produced through the action of biosynthetic enzymes encoded by select core genes residing in *epaA*–*epaR* (17, 34, 42). It is less clear how the genes in the variable region contribute to overall Epa composition. We discovered that the ability of phages to infect *E. faecalis* is mediated through Epa. Specifically, mutations in the core gene *epaR* and/or variable region genes including *epaS, epaW, epaX, epaY* and *epaAC* were sufficient to abrogate phage infection. Recently, it was discovered that mutation of *epaR* and *epaX* in *E. faecalis* prevented phage infection by excluding phage adsorption (16, 18). Here, we found that modifications made by *epa* variable genes are important for initial phage adsorption, although adsorption does not always lead to successful phage infection.

Phage infection occurs in three distinct stages: phage adsorption, host receptor engagement and phage DNA replication. Considering the phages from our study adsorb to non-susceptible bacterial strains, it is likely that non-cognate hosts either have an incompatible cognate receptor, lack the required phage receptor, or abort phage DNA replication. It is curious that all 32 phage resistant isolates reported in this study harbored mutations only in *epa* genes. This observation, combined with the knowledge that the phages adsorb to non-cognate host strains, indicates that *E. faecalis* preferentially subverts phage infection by mutating *epa*. We hypothesize that another factor is required for productive phage infection. This second factor is likely a bonafide phage receptor that facilitates DNA entry and may be an essential protein, as only *epa* mutations arose in phage resistant *E. faecalis* isolates. We propose that Epa is the attachment factor that positions phages in proximity of an unidentified receptor required for DNA ejection.

Previous studies have demonstrated that inactivation of core *epa* genes in *E. faecalis* affect bacterial fitness (16, 17). However, we have a limited understanding of the contribution of *epa* variable genes in this context. Similar to a recent report that an *epaX* mutant strain of *E. faecalis* has an intestinal colonization defect (34), we demonstrated that an *epaS* mutant strain is also impaired in intestinal colonization. Importantly, we show that *epaS* is a colonization determinant within the context of an unperturbed microbiota. This shows that Epa cell surface decorations help *E. faecalis* compete in a complex microbial community. In addition, an *epaS* mutant was impaired in its ability to transfer from mother to infant and establish productive colonization. Considering enterococci are life-long colonizers of humans and animals, this observation raises interesting questions about whether enterococci are transferred directly from mother to infant or if they are acquired from the environment following birth.

*E. faecalis* intestinal adaptation is facilitated by its inherent resistance to environmental stressors encountered in the mammalian gut such as low pH, high osmolarity and bile salts (16, 31, 43-45). Therefore, Epa likely plays a critical role in the survival of *E. faecalis* when encountering intestinal environmental stresses. *epa* genes aid in the ability of *E. faecalis* to tolerate environmental stress which is demonstrated by phage resistance and susceptibility to cell wall targeting antibiotics upon loss of functional *epa* genes. When challenged with vancomycin, an *epaS* mutant of *E. faecalis* in the mouse intestine lacks the ability to efficiently expand its population when bacterial diversity is diminished. These data suggest that the *epaS* mutant cannot tolerate vancomycin selection and remains a minority member of the microbiota.

In conclusion, we believe that phages like those described in this study are candidates for the development of straightforward therapeutics that could be used in conjunction with current antibiotic therapies to curtail the overgrowth of multidrug-resistant enterococci in vulnerable patients. We also believe that these data emphasize the importance of understanding phage infection mechanisms for the future development of phage cocktails. This information could help reduce the risk of developing phage resistance during therapy.

## Materials and Methods

### Bacteria and bacteriophages

A list of the bacterial and bacteriophage strains used in this study can be found in Table S3. *E. faecalis and E. faecium* were grown with aeration on brain heart infusion (BHI) broth or on BHI agar at 37°C. *Escherichia coli* was grown on Lennox L broth (LB) with aeration or on LB agar at 37°C. When necessary, for the selection of *E. coli* or *E. faecalis*, 15 µg/ml chloramphenicol (Research Products International) was added to the media. Growth conditions for the generation of mutant strains of *E. faecalis* by allelic exchange were as described by Thurlow et al. (46). Phage sensitivity assays were performed on Todd-Hewitt broth (THB) agar. The library of 19 enterococcal specific bacteriophages were obtained through the Biological Defense Research Directorate of the Naval Medical Research Center (NMRC).

### Determination of phage host range

The lytic activities of the 19 phages from the NMRC collection were screened against 21 different *E. faecalis* strains using a standard spot assay (27, 28). 250 µl of a 1:5 dilution of an overnight (O/N) culture of *E. faecalis* was mixed with 5 ml of THB top agar (0.35% agar) and poured onto the surface of a THB agar plate (1.5% agar). Both top agar and base agar were supplemented with 10 mM MgSO_4_. 5 µl of each phage lysate was spotted on the bacterial overlay plate. The plates were incubated at 37°C O/N, and *E. faecalis* sensitivity to individual phages was indicated by either clear, opaque or no clearing spots which indicated infection, weak infection and no infection, respectively.

### Isolation of phage resistant *E. faecalis strains*

250 µl of a 1:5 dilution of an O/N culture of host bacteria was mixed with 10 µl of serially diluted phage and 5 ml of pre-warmed THB top agar. Phage-bacterial mixtures were poured onto the surface of THB agar plates. The plates were incubated at 37°C until phage-resistant colonies appeared in the zones of clearing. The presumptive resistant colonies were passaged four times by streaking single colonies onto BHI agar. The phage-resistant phenotypes were confirmed by spot assays (27, 28).

### Phage adsorption assay

An O/N bacterial culture was pelleted at 3220 x *g* for 10 minutes and resuspended to 10^8^ cfu/ml in SM-plus buffer (100 mM NaCl, 50 mM Tris-HCl, 8 mM MgSO_4_, 5 mM CaCl_2_ [pH 7.4]). The cell suspensions were mixed with phages at a multiplicity of infection of 0.1 and incubated at room-temperature without agitation for 10 minutes. The bacteria-phage suspensions were centrifuged at 24,000 x *g* for 1 minute and the supernatant was collected to determine the phage concentration by plaque assay. SM-plus buffer with phage only (no bacteria) served as a control. Percent adsorption was determined as follows:

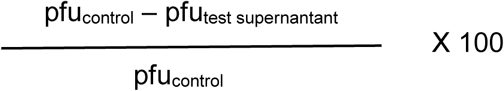

### Antibiotic susceptibility assay

*E. faecalis* was added to 7 ml of BHI broth (5×10^5^ cfu/ml final density). Daptomycin (Tokyo Chemical Industry) or vancomycin (Alvogen) were added to obtain the desired concentrations indicated in Figure 6. Cultures containing daptomycin were supplemented with 50 mg/ml CaCl_2_. Cultures were incubated at 37°C with aeration O/N. To determine viable cfu after O/N growth, cultures were serially diluted in phosphate buffered saline (PBS) and 10 µl were spotted onto BHI agar and incubated at 37°C O/N. Viable cfu/ml were determined by colony counting.

### Animals

C57BL6/J (conventional and germ free) male and female mice were used for these studies. For detailed information on specific animal experiments see the Supplementary Materials and Methods. All animal protocols were approved by the Institutional Animal Care and Use Committee of the University of Colorado School of Medicine (protocol number 00253).

### Data Availability

The DNA sequencing reads associated with this study are deposited at the European Nucleotide Archive (http://www.ebi.ac.uk/ena) under accession number PRJEB30526.

## Acknowledgements

We thank Biswajit Biswas for providing the NMRC phage collection. We thank Michael Gilmore for providing *E. faecalis* strain SF28073. This work was supported by National Institutes of Health R01 AI141479 (B.A.D.), K01 DK102436 (B.A.D.), R01 AI116610 (K.L.P.) and a National Institutes of Health and Food and Drug administration Inter-Agency Agreement AAI15010-001 (P.E.C.Jr.).

---

**Figure S1.** The cell surface of *E. faecalis epa* mutant strains is more positively charged. Cytochrome *c* binding to the surface of various *E. faecalis* strains indicates that mutations in the *epa* variable genes *epaS* and *epaAC* alter the net charge of the cell wall. -E (empty vector) and -C (complemented). *p<0.02, **p<0.008, ***p<0.001 by multiple comparisons one-way ANOVA and Tukey’s multiple comparison post-hoc test.

**Figure S2.** Mothers are colonized long-term by wild type and the *epaS* mutant strains of *E. faecalis*. Intestinal colonization levels of wild type *E. faecalis* V583 and the *epaS* mutant strain BDU61 from the mothers bearing pups used in the transmission experiments shown in Figure 8. Arrowheads indicate the day that the mothers gave birth to the pups.

**Table S1.** Spontaneous mutations in the *epa* cluster result in phage resistance in vitro.

**Table S2.** Spontaneous mutations in the *epa* cluster result in phage resistance in vivo.

**Table S3.** Bacterial strains, phages, plasmids and primers.

